# Tetraspanin 4 mediates migrasome formation via a two-stage mechanism

**DOI:** 10.1101/2022.07.11.499540

**Authors:** Raviv Dharan, Yuwei Huang, Sudheer Kumar Cheppali, Shahar Goren, Petr Shendrik, Michael M. Kozlov, Li Yu, Raya Sorkin

**Affiliations:** School of Chemistry, Raymond & Beverly Sackler Faculty of Exact Sciences, Tel Aviv University, Israel; Center for Physics and Chemistry of Living Systems, Tel Aviv University, Tel Aviv, Israel; The State Key Laboratory of Membrane Biology, Tsinghua University-Peking University Joint Center for Life Sciences, School of Life Sciences, Tsinghua University, Beijing, China; Beijing Frontier Research Center for Biological Structure, Beijing, China; School of Mechanical Engineering, The Ivy and Aladar Fleischman faculty of engineering, Tel Aviv University; Department of Physiology and Pharmacology, Sackler Faculty of Medicine, Tel Aviv University, Tel Aviv, Israel

**Author notes:** These authors contributed equally (Raviv Dharan, Yuwei Huang).

## Abstract

Migrasomes are recently discovered signalling organelles, enriched with tetraspanin proteins (TSPAN)^1^. They form by local swelling of retraction fibers, the cylindrical protrusions of cell membranes that form as a result of cell migration along external substrates. Migrasomes can grow up to several micrometers in diameter^2^, and allow cells to release contents such as chemokines at specific locations, hence, transmitting signals to surrounding cells through the relevant chemokine receptors. Recently, evidence emerged showing that migrasomes play essential roles in fundamental cellular processes such transfer of mRNA and proteins^3^, organ morphogenesis^4^, and mitochondria quality control^5^. Thus, understanding the mechanism of migrasome biogenesis is of outstanding importance. Previously, it was established that the molecules crucial for migrasome formation are tetraspanin proteins and cholesterol forming macrodomains in the migrasome membrane, while the physical forces driving local swelling of the retraction fibers originate from membrane tension and bending rigidity^1^. Yet, it remained unknown how and in which time sequence these factors are involved in migrasome nucleation, growth, and stabilization, and what are the possible intermediate stages of migrasome biogenesis.

---

Here we reveal that tetraspanin 4 (TSPAN4) mediates a two-stage process of migrasome formation. At the first stage, membrane swellings form along retraction fibers. At the second stage, TSPAN4 migrates toward and onto these swellings, which grow to migrasomes of several microns in size. This TSPAN4 recruitment to the swellings is essential for migrasome growth and stabilization. We demonstrate this mechanism by imaging migrasome generation in live cells and validate it by recreating the conditions leading to migrasome-like vesicle generation in a biomimetic model system. Based on these findings we propose that the major role of TSPAN proteins is in stabilizing the migrasome structure while the migrasome nucleation and initial growth stages can be driven by membrane mechanical stresses.

We followed the formation of migrasomes of Normal rat kidney (NRK) cells over-expressing TSPAN4-GFP stained with FM4-64 membrane dye under confocal microscopy. Migrasomes formed along retraction fibres following cell migration (Fig.1A). Closer to the cell body, FM4-64 enriched puncta could be seen along the retraction fibres (Fig.1A and supplementary movie1). Further from the cell, large migrasomes were observed, enriched with both Tspan4-GFP and FM4-64 (Fig.1A and supplementary movie1). The fact that mature migrasomes were enriched with TSPAN4 while the newly generated ones were not, suggests the initial stage of migrasome biogenesis to occur ahead of the TSPAN4 recruitment. To further test this hypothesis, we conducted time-lapse imaging of living cells by using structural illumination microscopy (SIM) (Fig. 1B). At an early stage, FM4-64 was locally enriched as puncta on the retraction fibers. The small puncta could move along the retraction fibers and coalesce to bulge out from the thin retraction fibers as small swellings. At this initial stage, Tspan4-GFP was relatively homogeneously distributed along the retraction fibers. At the next stage TSPAN4-GFP was gradually recruited onto the swellings, which grew into migrasomes (Fig. 1B, C). To address this process quantitatively, we followed the formation of 252 individual migrasomes. We divided the migrasomes into four groups based on the lipid and TSPAN4-GFP fluorescence intensity at the initial stage of migrasome formation versus the migrasome growth stage (detailed examples of these stages are shown in figure S1). The groups are defined as follows: (1) initial-stage is red (enriched in FM4-64), growth stage is yellow (enriched in FM4-64 and TSPAN4-GFP), (2) initial stage is red, growth stage is red, (3) initial stage is green (enriched in TSPAN4-GFP), growth stage is green, (4) initial stage is yellow, growth stage is yellow (Fig. 1D). Most of the mature migrasomes (almost 80% (Fig. 1D)) started to form from FM4-64 enriched small puncta and then grew in size concomitantly with further TSPAN4-GFP enrichment.

**Figure 1.**
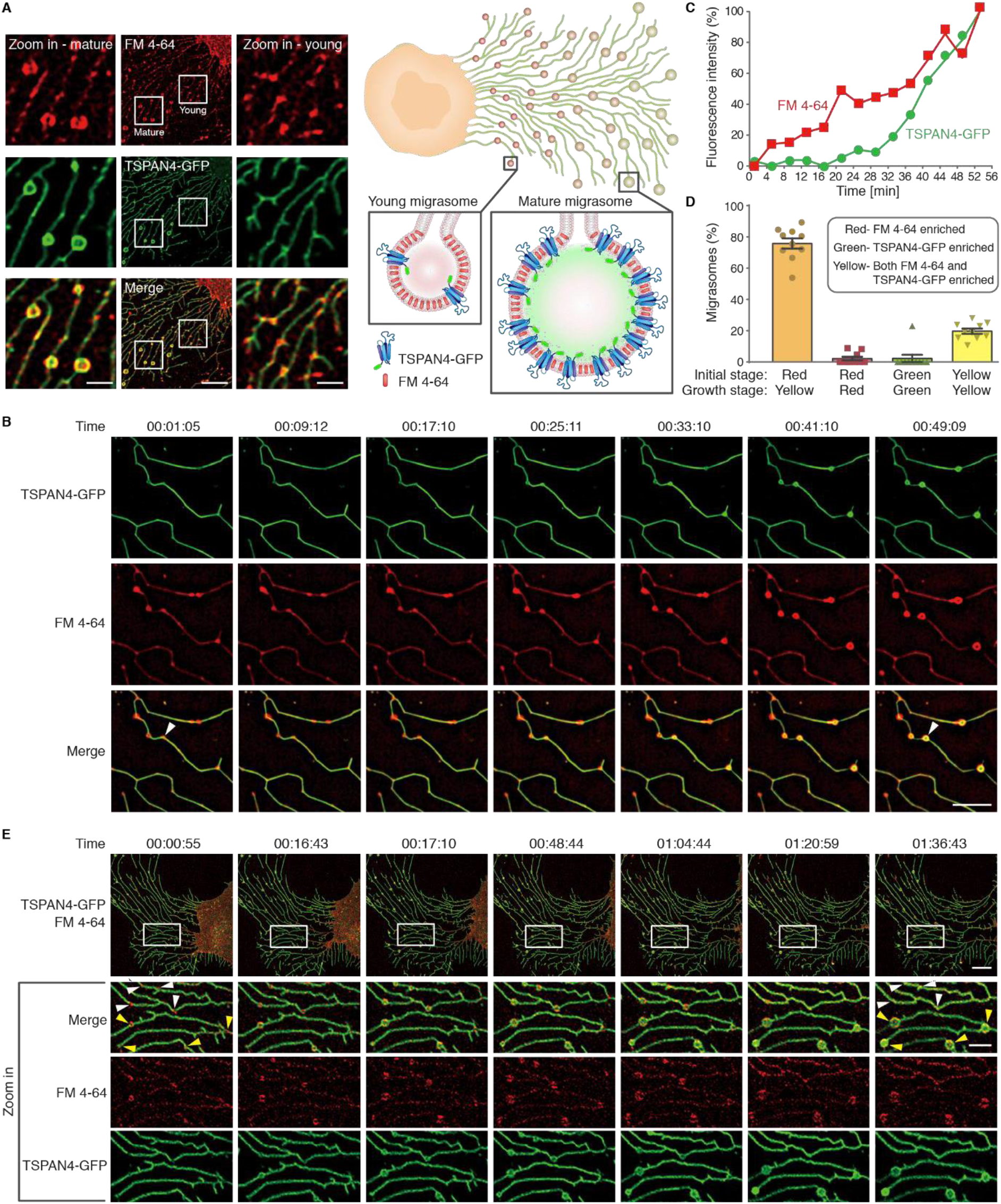
Two-stage mechanism of migrasome formation. (A) Confocal images of NRK TSPAN4-GFP (green) cells stained by FM4-64 (red). Scale bar, 10 μm; zoom in, 2.5 μm. On the right, schematic representation of young migrasomes with low TSPAN4-GFP concentration and mature migrasomes with TSPAN4-GFP enrichment. (B) Time-lapse images of NRK TSPAN4-GFP cells stained by FM4-64. Imaging by structural illumination microscopy (SIM). Scale bar, 5 μm. (C) Normalized fluorescence intensity as function of the time of TSPAN4-GFP and FM4-64 on a representative migrasome in B, indicated by a white arrow. Normalization based on retraction fiber fluorescence. (D) Statistical analysis of 4 different kinds of migrasomes during biogenesis (red-yellow, red-red, green-green, yellow-yellow), based on a series of time-lapse images of NRK TSPAN4-GFP cells stained by FM4-64 under confocal microscopy. N=252, from 11 individual cells from three independent experiments. (E) Confocal Time-lapse images of NRK TSPAN4-GFP cells stained by FM4-64. White arrows point to migrasomes that form and shrink back; yellow arrow heads point to growing migrasomes. Scale bar, 10 μm; zoom in, 3 μm. Time in B and E is hh:mm:ss.

We have further concluded that TSPAN4 recruitment to the migrasomes is necessary for migrasome stabilization. We observed two populations of migrasomes (Fig. 1E). The population indicated by the white arrows initially increased in size, but eventually shrunk back within the time course of the experiment. The migrasomes indicated by yellow arrows grew and stably maintained their large sizes throughout the experiment. The main visible difference between these two types of migrasomes was in their TSPAN4 signal which was increased in the stable migrasomes. Overall, these results strongly suggest a two-stage mechanism of migrasome biogenesis, the first stage being formation of FM4-64 positive small swellings, and the second stage being migrasome growth and stabilization through TSPAN4 recruitment.

In order to get insight into the main factors underlying the two-stage mechanism of migrasome biogenesis, we designed a biomimetic system emulating a cell with a retraction fiber and enabling an experimental simulation of migrasome formation. We used optical tweezers combined with confocal fluorescence microscopy and micropipette aspiration^6,7^. As a model for the cell, we used a giant plasma membrane vesicle (GPMV)^8^, which we generated from HEK293T cells expressing TSPAN4-GFP (Fig. 2). To imitate a retraction fiber, we pulled a membrane tube out of an aspirated GPMV by attaching a polystyrene bead to the vesicle and then moving the bead away by optical tweezers (Fig.3A). This setup enabled us to control the membrane tension of the GPMV and, hence, the tube diameter through setting the aspiration pressure, to measure the force pulling the tube by the optical tweezers, to perform confocal fluorescence imaging of the system, and monitor the system evolution in real time via bright-field microscopy.

**Figure 2.**
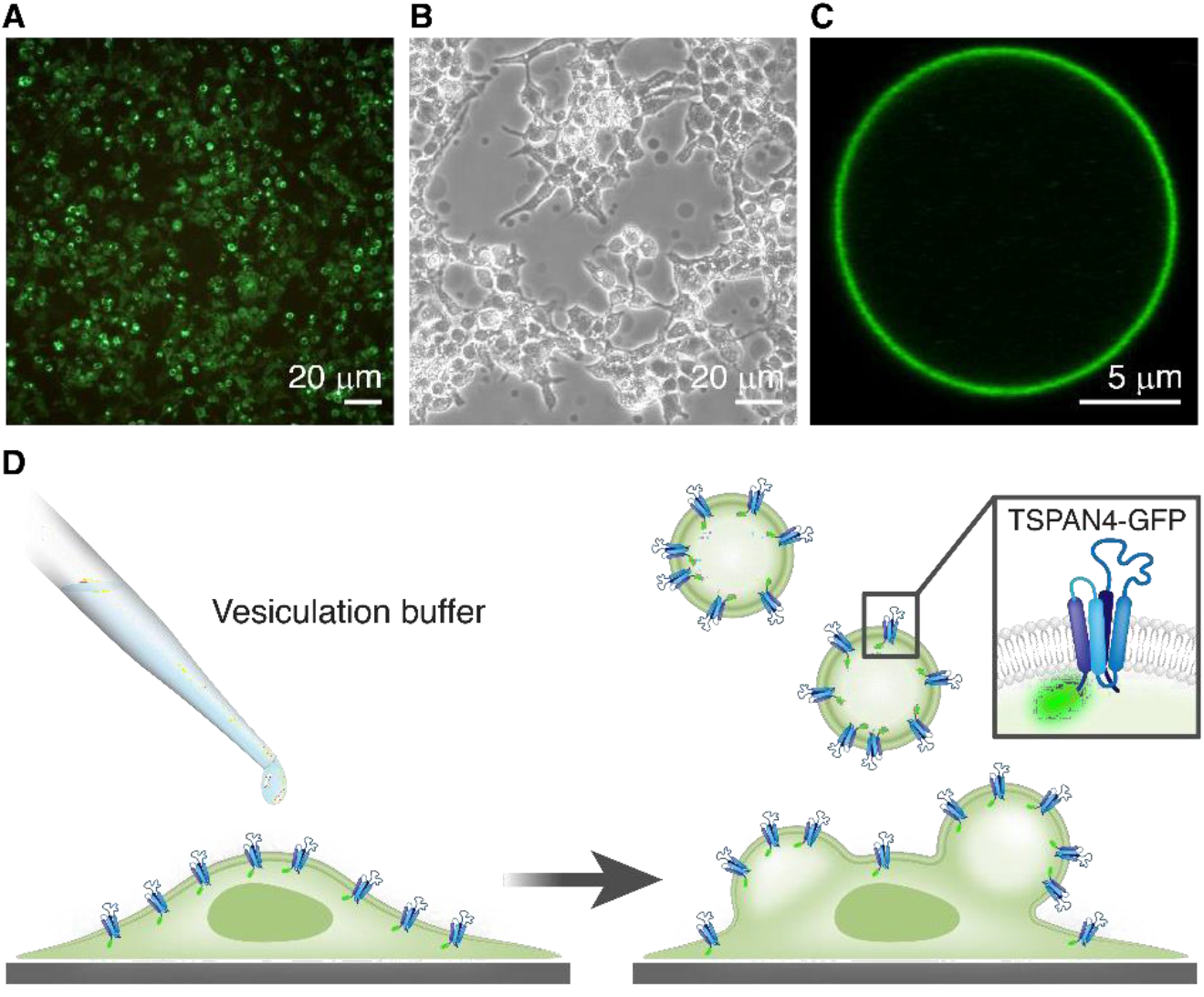
Formation of giant plasma-membrane-vesicles (GPMVs) from transfected HEK293T cells expressing TSPAN4-GFP. (A) Microscopy image of HEK293T cells expressing TSPAN4-GFP 24 hours after transfection. (B) Microscopy image of HEK293T cells, expressing TSPAN4-GFP, after treatment with a vesiculation buffer. GPMVs, which appear dark in phase contrast image, can be seen floating in the sample or attached to the cells. (C) Confocal microscopy images of GPMVs containing TSPAN4-GFP. (D) Schematic representation of a transfected cell, treated with vesiculation buffer, producing GPMVs with TSPAN4-GFP in their membrane.

To simulate the initial stage of the migrasome formation we reasoned that the generic factor driving local swellings of membrane tubules might be an abrupt increase of membrane tension, which is known to lead to tube pearling instability^9^. A crucial role of membrane tension in migrasome formation is supported by the previous work^1^. To test this idea, we designed an experiment of a two-step tension application to a membrane tube. First, the tube was pulled out of a GPMV subject to a relatively low membrane tension such that the tube radius was relatively large. Next, we rapidly increased the GPMV aspiration pressure and, hence the membrane tension^10^ (supplementary movie2). The time of the tension increase was substantially shorter than the time needed for the tube relaxation to a new equilibrium configuration of a homogeneous cylinder with a reduced cross-sectional radius corresponding to the new level of the tension, which required a slow decrease of the intra-tubular volume through liquid flow into the GPMV. This condition of a transiently constant volume corresponded to that of the pearling instability^9^. Indeed, the abrupt increase of tension led to generation of the migrasome-like local swellings of the tube (Fig. 3A, S2). The swellings were able to move along the tube (Figure S3). The constricted regions of the tube were enriched with TSPAN4 whereas the swellings were enriched with DiI-C12 (Figure 3B), which is a membrane dye known to preferentially partition in disordered lipid phase^11^.

**Figure 3.**
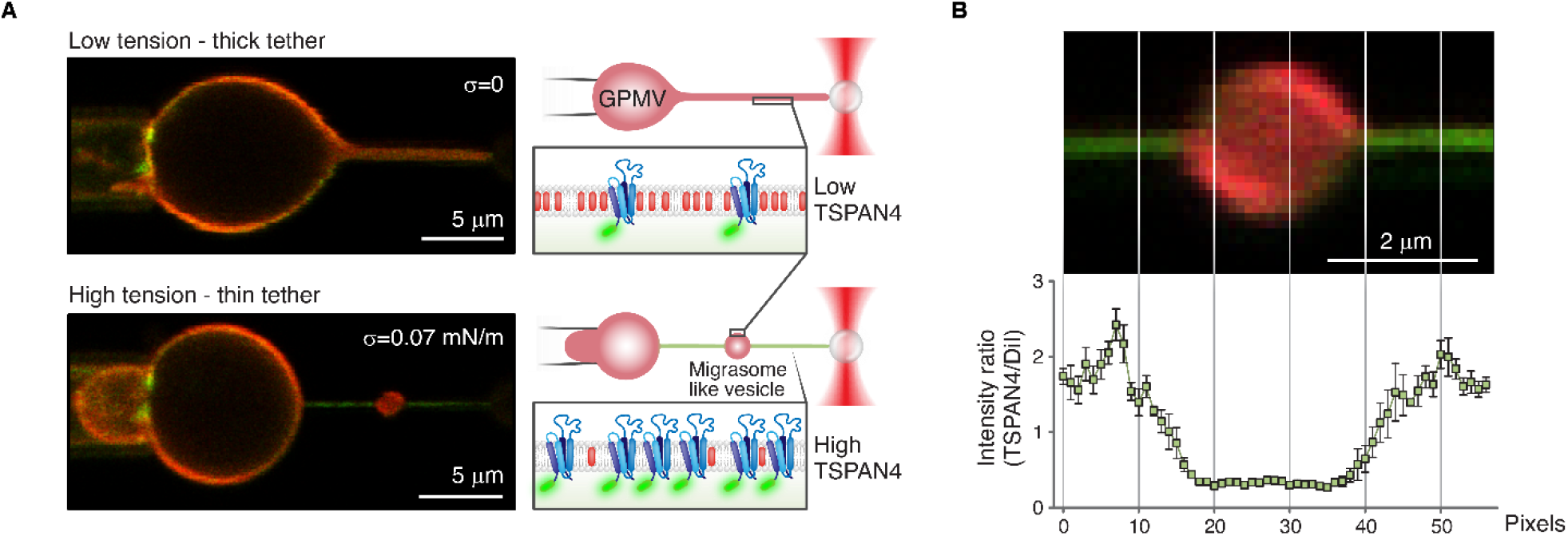
Swelling formation on membrane tube induced by rapid tension increase. (A) Confocal microscopy images of a membrane tube pulled from GPMVs containing TSPAN4-GFP (green) and DiI-C12 (red) aspirated with micropipette. In the top image the suction pressure was zero (corresponds to zero tension applied, T=0). Next, the tension increased immediately to 0.07 mN/m. On the right side, schematic representation of tension-induced swelling formation assay. (B) TSPAN4-GFP and DiI-C12 fluorescence intensity ratio of membrane tubes containing a swelling (n=4 membrane tubes, having swelling with relatively same size, pulled from 4 vesicles, error bars are SEM). On the right, a representative image of a membrane tube containing a swelling.

As a following step, we examined whether TSPAN4 in our model self-organizes into clusters. The functionality of tetraspanins is thought to be dependent on their ability to associate among themselves and with other integral proteins and adhesion molecules, forming a distinct class of membrane domains^12,13^. TSPAN domains have been shown to be dynamic and varying in composition and organization between different cell lines and cell states^14^. In a previous study, TSPAN4 molecules were found to organize into highly dynamic clusters on the retraction fibers^1^. We found that in our system, while being highly enriched in the tubular membranes, TSPAN4 was ununiformly distributed along the tubes and formed mobile puncta (Fig. S4A), which can be classified as clusters, as previously observed in cellular retraction fibers^1^. We further demonstrated that TSPAN4 clusters formed also on the flat membrane of GPMVs as a result to shear forces induced by buffer flow (Figure S4B). In this experiment, the GPMVs were injected into a microfluidics chamber under high pressure, which led to substantial shear forces. Altogether our results demonstrate the tendency of TSPAN4 to cluster and form domains.

Next, we sought to examine whether our model system is able to recreate TSPAN enrichment in the swellings, as observed for the migrasome biogenesis in live cells. In 7 out of 23 experiments, we observed TSPAN4-GFP migration to the swellings prior to tube rupture (Fig. 4A). TSPAN4 migrated to the swellings in the form of TSPAN-enriched domains and stayed on the swellings until the tube rupture. To quantify the swelling enrichment with TSPAN4, we calculated the relative intensities of TSPAN4-GFP and DiI-C12 on the swellings compared to the GMPV, which increased with time (Fig. 4B). Furthermore, TSPAN4 containing tubes exhibited an augmented tendency to rupture as compared to the control tubules i.e., without TSPAN4 (Fig.S5A). The swellings were observed to remain intact after the rupture (Fig. S5B). The likely reason for the lack of partitioning of TSPAN4 domains to the swellings in all the experiments was the tubule rupture, which did not leave enough time for the domains to migrate. The rupture may be promoted by the membrane structural defects emerging along the boundaries of the TSPAN4 enriched domains as suggested by previous reports showing that the lipid phase separation can lead to the tubule rupture^15^.

**Figure 4.**
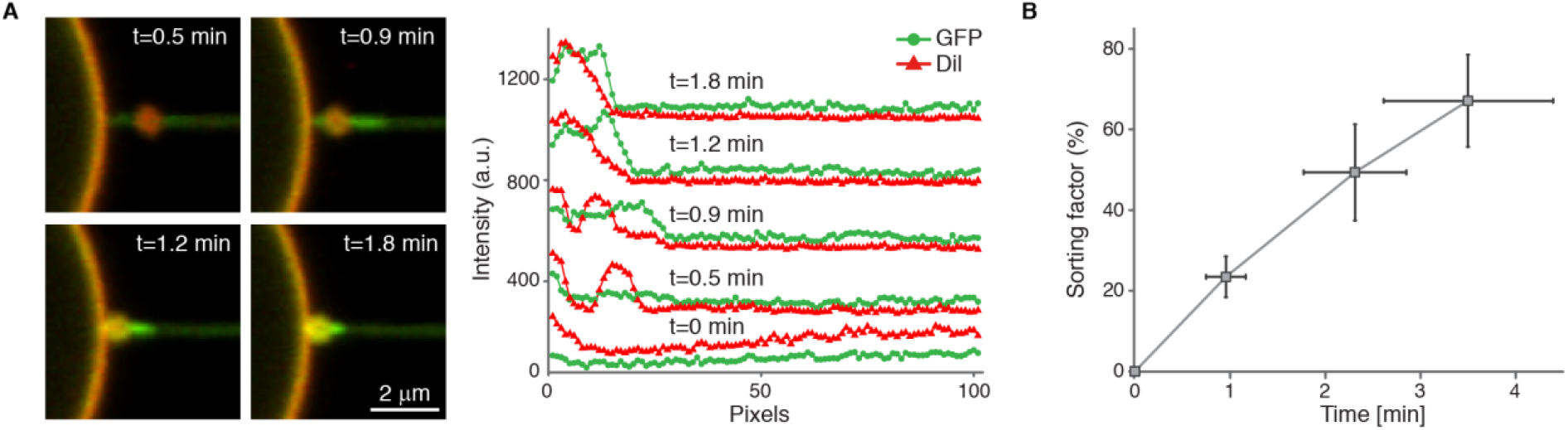
TSPAN4 domains recruitment to the tubule swellings. (A) Time-lapse confocal microscopy images of a membrane tube pulled from a GPMV containing TSPAN4-GFP (green) and DiI-C12 (red). Following swelling formation, TSPN4 clustering followed by TSPN4 swelling enrichment was observed. On the right the fluorescence intensity of DiI-C12 and TSPAN4-GFP along the tube and the swelling at the indicated times (t=0 correspond to the intensity of the tube just before the swelling formation). (B) Percentage change of TSPN4 sorting factor, 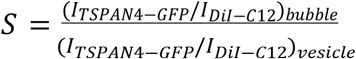, which is the fluorescence intensity ratio of GFP and DiI-C12 on the swelling compared to the vesicle, as function of time (n= 7 membrane tubes pulled from 6 vesicles, error bars are SEM).

We suggest the following explanation for TSPAN4 recruitment to migrasomes. While TSPAN4 molecules exhibit high positive intrinsic curvature corresponding to an effective molecular shape of an inverted cone (we characterize this in detail in a separate publication), TSPAN4 assembly into clusters and larger domains can reduce its intrinsic curvature. This leads to migration of the domains onto the membrane swellings that have a smaller curvature and, therefore, a better curvature compatibility with the domains. The hypothesis of lower intrinsic curvature of large TSPAN-enriched domains compared with single TSPAN proteins is further supported by the finding that TSPANs associate with cholesterol^1^. The cholesterol molecules in mixtures with common lipids have a cone-like effective shape corresponding to a negative intrinsic curvature^16^. Because of the opposing geometries, a complex of cholesterol and TSPAN must have an intrinsic curvature substantially lower than that of individual TSPAN molecules.

Finally, we tested whether our model system exhibits slower dissipation of membrane swellings in the presence of TSPAN4. We conducted control experiments with GPMVs that did not contain over-expressed TSPAN4. Strikingly, the swellings demonstrated a behavior very similar to that of the TSPAN4-depleted migrasomes in live cells, where they dissipated in the absence of TSPAN4 (Fig. 5). Specifically, after an abrupt tension increase in a GMPVs lacking TSPAN4, the swellings formed along the membrane tube, similarly to the results presented in Figure 3, and then rapidly disappeared within on average a 30 second time span after formation (Fig.S5C). In the experiments with GMPVs containing TSPAN4, however, the formed swellings were much more stable (Fig. 5), their lifetime being at least 5 times longer than that measured in the control experiments (Fig.S5C)).

**Figure 5.**
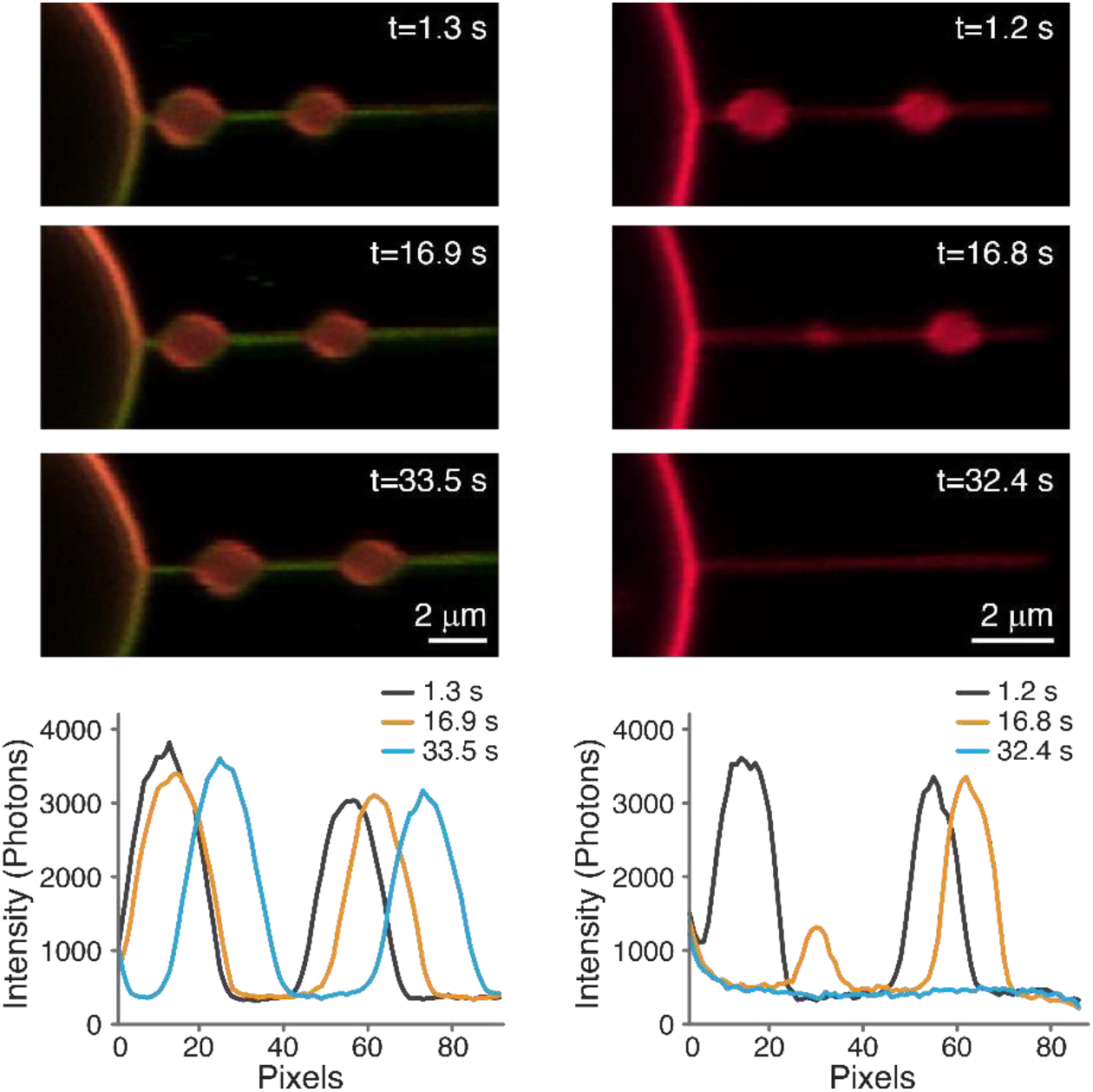
TSPAN4 inhibit swelling dissipation. Time-lapse confocal microscopy images of membrane tubes with a swelling, dyed with DiI-C12 (red) in the presence of (left images) and without (right images) TSPAN4-GFP (green). At the bottom: fluorescence intensity profiles of DiI-C12 along the membrane tube and swellings at the indicated times.

Overall, we investigated the temporal pathway of migrasome formation in conjunction with TSPAN4 dynamics on retraction fibers of live cells and designed a bio-mimetic system emulating these processes in a minimal artificial system. In this system, we recreated the crucial aspects of migrasome biogenesis observed in live cells: formation on membrane tubes of initial migrasome-like local swellings having relatively low TSPAN concentrations, formation of TSPAN domains and their recruitment to the swellings, dissipation of the swellings in the absence and their stabilization in the presence of TSPAN.

Based on the obtained results, it can be concluded that the migrasome formation proceeds in two sequential steps: formation of local swellings on the tubular retraction fibers, and stabilization of these swellings by TSPAN-based membrane domains. Our results suggest that the migrasome biogenesis can be driven by a very limited set of factors: the first step driven by membrane tension with possible involvement of additional factors in live cell membranes, and the second step controlled by specific proteins of the TSPAN family. The particular cellular mechanisms by which membrane tension and TSPAN-based clusters shape the migrasomes, while being partially addressed in the previous work^1^, await substantial elaboration. Our results illuminate the mechanism of migrasome biogenesis and demonstrate the crucial role of TSPAN proteins in membrane shaping processes.

## Acknowledgements

RS acknowledges support by the ISRAEL SCIENCE FOUNDATION (grant No. 1289/20). SKC acknowledges support by the Ratner Center for Single Molecule Science. MMK was supported by Deutsche Forschungsgemeinschaft (DFG) through SFB 958 “Scaffolding of Membranes”, and Israel Science Foundation grant 3292/19, and holds Joseph Klafter Chair in Biophysics. YH acknowledges support by the National Natural Science Foundation of China (32070691).

## Materials and methods

### Cell culture

HEK293T and Normal rat kidney cells were cultured at 37°C and 5% CO_2_ in DMEM supplemented with 10% serum and 1% penicillin-streptomycin.

### Cell Imaging

#### Cell sample preparation

NRK Tspan4-GFP cells were seeded into 3.5 cm glass bottom confocal dish which was pre-coated with 10 μg/mL fibronectin and grew for 15h. The cells were stained with 5 μg/mL FM4-64 for 15 minutes.

#### Confocal microscopy imaging

Cell imaging were conducted under galvanometer scanning mode using a NIKON A1 confocal microscope fitted with a 100× oil objective. The laser power was 0.5% for 488 nm and 3% for 561 nm and each field of 1024 × 1024 pixels was imaged. For time-lapse imaging, the interval was 4 minutes and the duration was ∼5 hours.

#### SIM imaging

The prepared cells were imaged by structured illumination microscopy (Nikon N-SIM S) with 4 minutes interval and 1.5 hours duration, and then reconstructed by a standard stack-reconstruction process.

### Quantitative analysis of migrasomes

252 migrasomes from 4 movies were analyzed, each movie 4 hours in length, with 60 frames in total. Fully formed migrasomes were chosen in the last frames of the movies, and then tracked back to their initial stage in the earlier frames. Only migrasomes that formed in the time course of the movie were included in the analysis. For the growing stage, several frames were included in the analysis in order to validate the presence of fully formed migrasomes due to possible changes in the focus. An example of initial stage, starting from red puncta, and growing stage is shown in supplementary figure 1.

### TSPAN4 Expression Plasmids, cell transfection and giant plasma membrane vesicles (GPMVs) isolation

Complementary DNAs of tetraspanin 4 were cloned into pEGFP-N_1_. HEK293T cells were plated in 25 cm^2^ flask coated with poly-L-lysine)Sigma) to keep the cells attached during the blebbing process and to minimize cell debris in solution. At 50% confluency, cells were transiently transfected with 5 μg DNA using Lipofectamine 2000 (Invitrogen) according to the manufacture’s protocols and then grown 24 hours for protein expression. GPMVs were produced according to a published protocol^17^. Briefly, following TSPAN4-GFP expression, for most of the experiments the cells were stained with DiI-C12 membrane dye (Invitrogen), washed with GPMV buffer (10 mM HEPES, 150 mM NaCl, 2 mM CaCl2, pH 7.4) twice, and incubated with 1 mL of GPMV buffer containing 1.9 mM DTT (Sigma) and 27.6 mM formaldehyde (Sigma). Secreted GPMVs were then collected and isolated from the cells and immediately used for the optical trapping experiments.

### Proteolysis and Mass Spectrometry Analysis

In order to confirm TSPAN4 overexpression in the GPMVs, we conducted mass spectrometry measurements which showed that TSPAN4 abundance in GPMVs generated by TSPAN4-transfected cells increased 1562-fold compared to control GPMVs (GPMVs that were generated from normal HEK293 cells as described above). The samples were brought to 10mM DTT, 100 mM Tris and 5% SDS, boiled in 95 °C for 10 minutes and sonicated twice for 10 minutes (5’, 10-10, 90%). The samples were precipitated in 80% acetone overnight and washed 3 times with 80% acetone. The protein pellets were dissolved in 8.5 M Urea and 400 mM ammonium bicarbonate. Protein amount was estimated using Bradford readings. Proteins reduced with 10 mM DTT (60 °C for 30 min), modified with 40 mM iodoacetamide in 100 mM ammonium bicarbonate (room temperature for 30 minutes in the dark) and digested in 1.5 M Urea, 66 mM ammonium bicarbonate with modified trypsin (Promega), overnight at 37 °C in a 1:50 (M/M) enzyme-to-substrate ratio. An additional trypsin digestion was performed for 4 hours at 37 °C in a 1:100 (M/M) enzyme-to-substrate ratio. The resulting tryptic peptides were desalted using C18 stage tips (homemade, 3M company, USA) dried and resuspended in 0.1% Formic acid. The peptides were resolved by reverse-phase chromatography on 0.075 × 300-mm fused silica capillaries (J&W) packed with Reprosil reversed phase material (Dr Maisch GmbH, Germany). The peptides were eluted with linear 60 minutes gradient of 5 to 28% 15 minutes gradient of 28 to 95% and 15 minutes at 95% acetonitrile with 0.1% formic acid in water at flow rates of 0.15 μl/min. Mass spectrometry was performed by Q Exactive Plus mass spectrometer (Thermo) in a positive mode using repetitively full MS scan followed by high collision dissociation (HCD) of the 10 most dominant ions selected from the first MS scan. The mass spectrometry data was analyzed using Proteome Discoverer 2.4 software with Sequest (Thermo) search algorithm against Human Uniprot database with 1% FDR. Semi quantitation was done by calculating the peak area of each peptide based its extracted ion currents (XICs), and the area of the protein is the average of the three most intense peptides from each protein.

### Tube pulling from aspirated GPMVs

The experiments were performed using a C-trap ® confocal fluorescence optical tweezers setup (LUMICKS) made of an inverted microscope based on a water-immersion objective (NA 1.2) together with a condenser top lens placed above the flow cell. The optical traps are generated by splitting a 10W 1064-nm laser into two orthogonally polarized, independently steerable optical traps. To steer the two traps, one coarse-positioning piezo stepper mirror and one accurate piezo mirror were used. Optical traps were used to capture polystyrene microbeads. The displacement of the trapped beads from the center of the trap was measured and converted into a force signal by back-focal plane interferometry of the condenser lens using two position-sensitive detectors. The samples were illuminated by a bright field 850-nm LED and imaged in transmission onto a metal-oxide semiconductor (CMOS) camera. **Confocal fluorescence microscopy:** The C-Trap uses a 3 color, fiber-coupled laser with wavelengths 488, 561 and 638 nm for fluorescence excitation. Scanning was done using a fast tip/tilt piezo mirror. For confocal detection, the emitted fluorescence was descanned, separated from the excitation by a dichroic mirror, and filtered using an emission filters (Blue: 500-550 nm, Green: 575-625 nm and Red: 650-750 nm). Photons were counted using fiber-coupled single-photon counting modules. The multimode fibers serve as pinholes providing background rejection.

#### Experimental chamber

PDMS walls were placed on the bottom cover slip (Bar Naor) and mounted onto an automated XY-stage. The GPMVs sample was added to the chamber and after about 15 minutes, a few drops of oil were put on the sample surface to prevent evaporation. A micropipette aspiration setup including micromanipulator (Sensapex) holding a micropipette with diameter of 5 μm (Biological industries) connected to a Fluigent EZ-25 pump was integrated to our optical tweezers instrument. Before each experiment, the zero-suction pressure was found by aspirating a 3.43 μm polystyrene bead (Spherotech) into the pipette and reducing the suction pressure until the bead stopped moving. A membrane tube was pulled from aspirated GPMVs using beads trapped by the optical tweezers. First, a membrane tube was pulled at relatively low suction pressure (0.05-0.1 mbar, correspond to 1.2-2×10^−5^ N/m membrane tension), then the suction pressure was reduced to the zero (corresponds to zero applied membrane tension) for about 15 seconds. Then, we increased instantaneously the suction pressure to values in the range of 0.2-0.7 mbar (correspond to 4-10×10^−5^ N/m membrane tension). For confocal imaging the 488 nm and 532 nm lasers were used for GFP and DiI-C12 excitation with emission detected in three channels (Blue, Green, Red). To induce shear forces on the GPMVs, the GPMVs were injected into a 5-channel laminar flow cell (LUMICKS, Amsterdam, the Netherlands).

### Data Analysis

Data acquisition was carried out using Bluelake, a commercial software from Lumicks. This software stores experimental data acquired during experiments with the C-trap in HDF5 files, which can be processed using Lumicks’ Pylake python package. Images of the confocal scans were reconstituted from photon count per pixel data in the HDF5 files using Pylake. All data analysis was performed with custom-written Python scripts. Fluorescence intensity profiles were obtained from the images by averaging the photon count of the relevant fluorescent channel (Blue or Green) in the region of interest.

