## Supplementary information for "Tetraspanin 4 mediates migrasome formation via a two-stage mechanism"

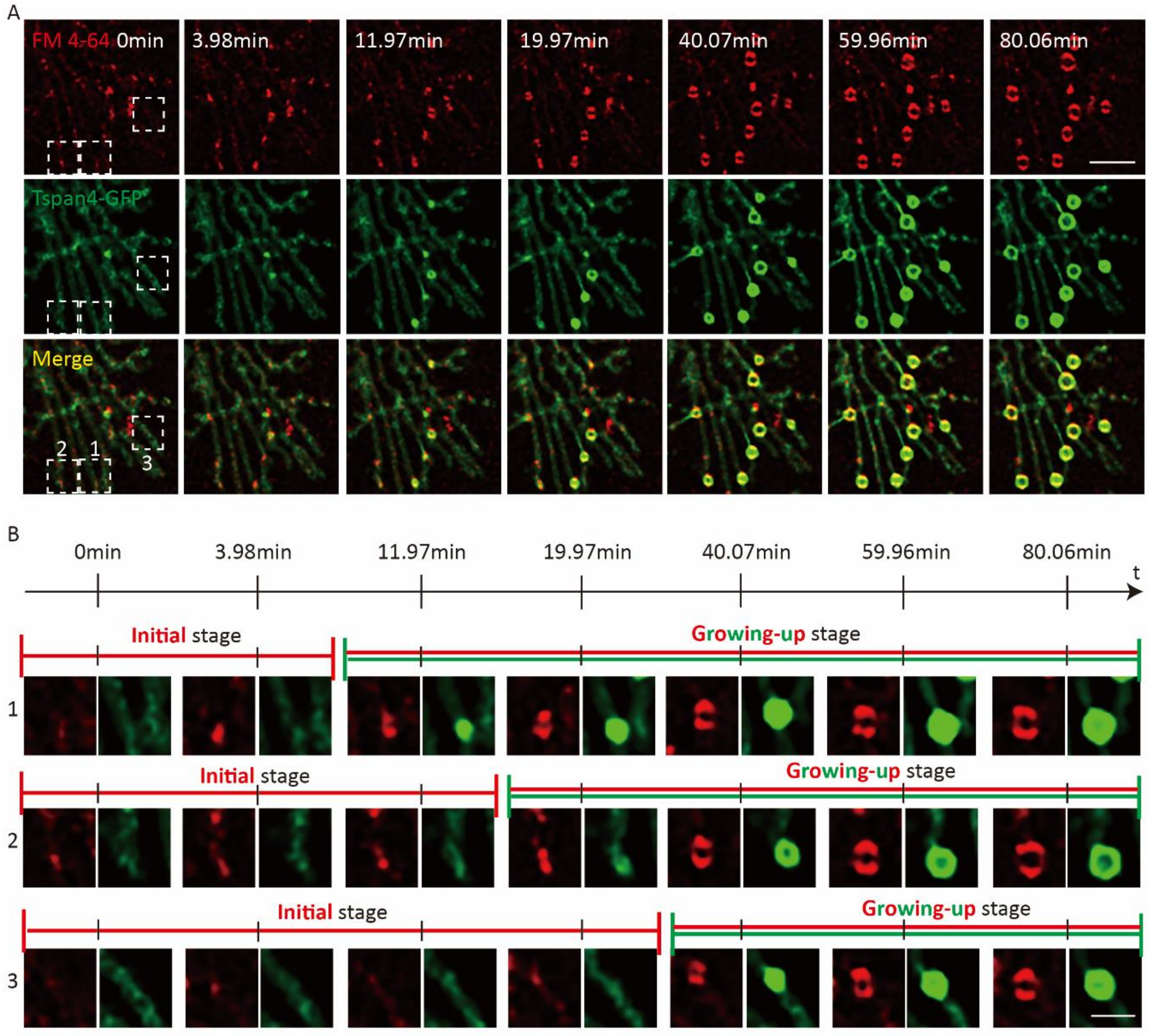


Figure S1. A. Confocal Time-lapse images of NRK Tspan4-GFP cells stained by FM4-64 (5μg/mL). Green, Tspan4-GFP; red, FM4-64. Scale bar, 5μm. B. Representative migrasome 1, 2 and 3 selected from A, zoomed in. The initial stage and growing stage are shown. Scale bar, 2μm.

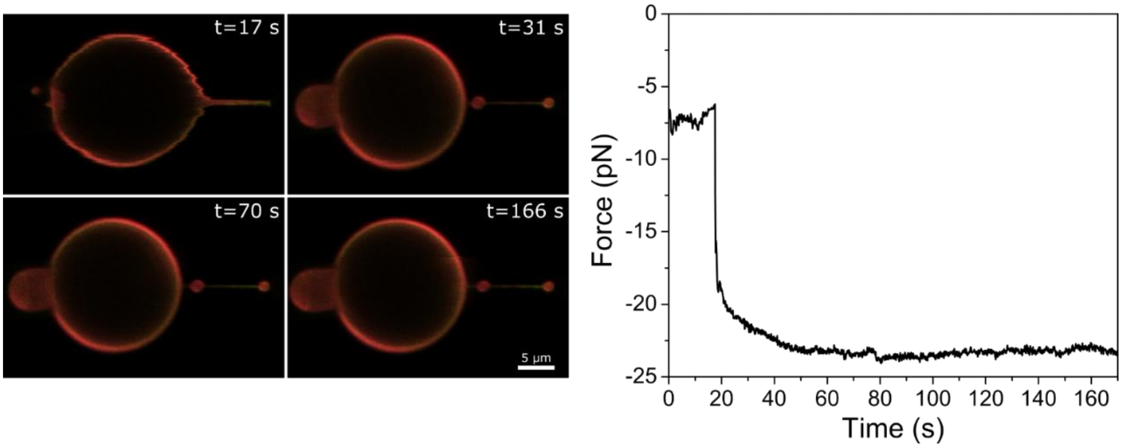


Figure S2. Swelling formation induced by tension increase. On the lest time-lapse confocal images of membrane tube pulled from GPMV containing TSPAN-GFP (green) and dyed with DiIC12 (red) aspirated at zero suction pressure (t=17 s). At t=20 s, the suction pressure was increased instantly to 0.25 mbar, which induce swellings formation on the tube. On the right, the force acting on the tube during the experiment.

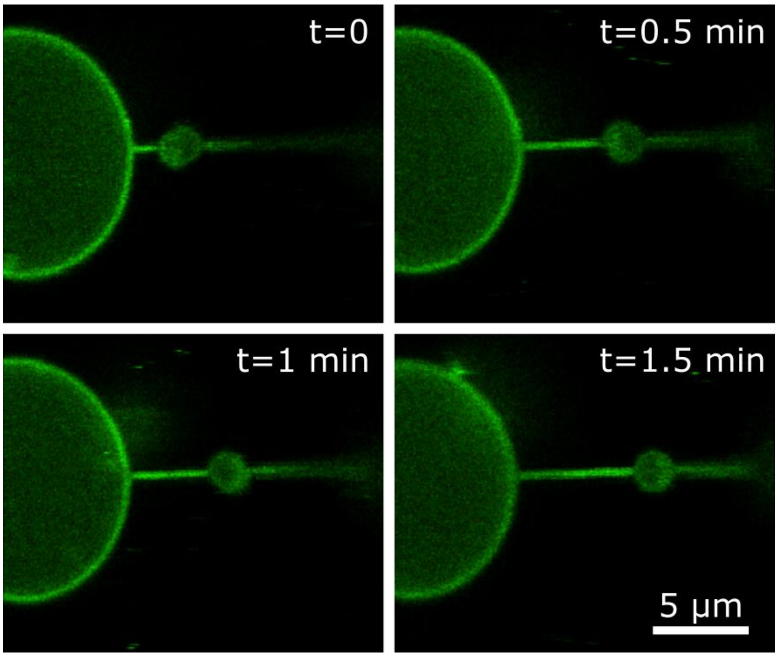

Figure S3. Swelling movement along membrane tube. Time-lapse confocal microscopy images of tube pulled from GPMV containing TSPAN4-GFP. After tension induced swelling formation, the swelling was able to move freely along the membrane tube.

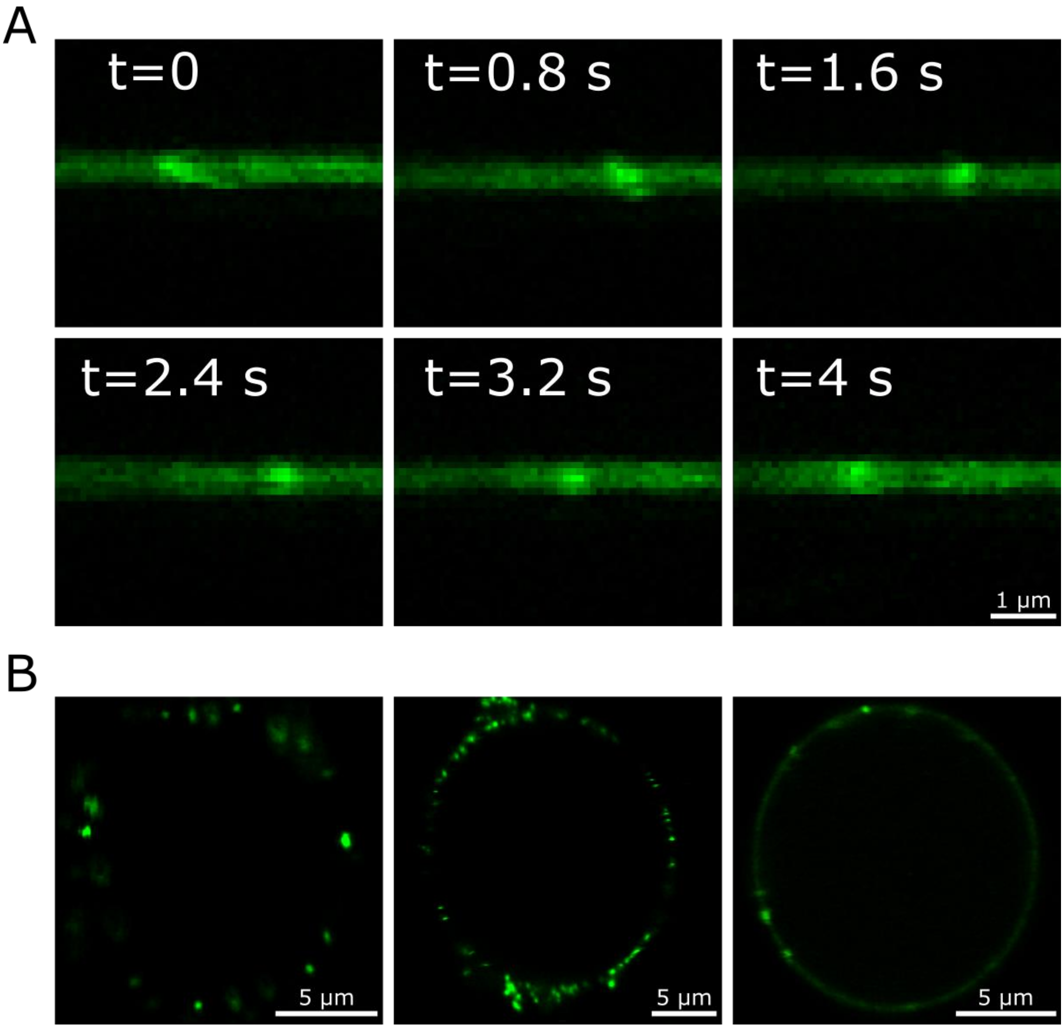


Figure S4. Formation of TSPAN4 domains induced by membrane curvature and shear forces. (A) Time-lapse confocal microscopy images of membrane tube enriched by TSPAN4-GFP at different time points as indicated. The first image (t=0) was taken 21 min after pulling the tube from giant plasma membrane vesicle (GPMV). GFP puncta, which relates to TSPAN4 domains can be seen on the membrane tube. (B) Confocal microscopy images of GPMVs, containing TSPAN-GFP, after flowing through microfluidic channel. The images show TSPAN4 domains on the GPNVs, which can induce significant membrane remodeling. Most of the vesicle were ruptured during the flow in the microfluidic channel.

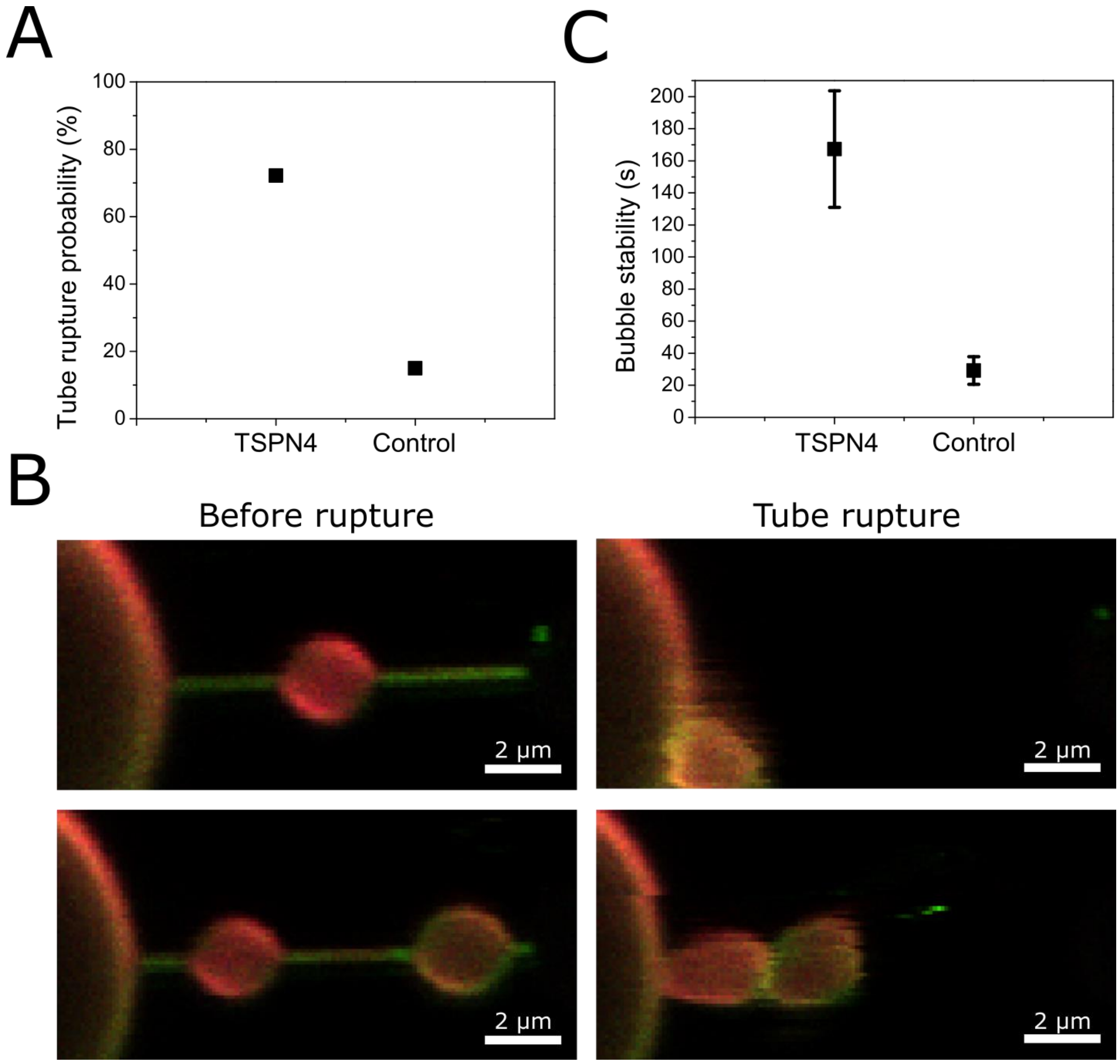


Figure S5. Swelling stability and tube rupture. (A) Percentage of tube rupture events with or without TSPAN4 (TSPAN4 n=18 membrane tubes pulled from 8 vesicles, control n=20 membrane tubes pulled from 8 vesicles). (B) Confocal microscopy images of swellings on a membrane tube pulled from GPMV containing TSPAN4-GFP (green) and DiI-C12 (red). On the left, the last frame before tube rupture, and on the right, the frame right after the tube rupture. The images show that the swellings were intact after the rupture. (C) Swelling stability on membrane tubes with or without TSPAN4. In the presence of TSPAN4 in most of the experiments the swelling was stable, and the measurements was ended due to tube rupture. Hence, the stability of the swelling in the presence of TSPAN4 is probably higher than indicated (TSPAN4 n=23 swellings on 18 membrane tubes pulled from 8 vesicles, Control n=24 swellings on 20 membrane tubes pulled from 8 vesicles. (Error bars are SEM)
